# Encoding of Dynamic Facial Information in the Middle Dorsal Face Area

**DOI:** 10.1101/2022.07.24.501335

**Authors:** Zetian Yang, Winrich A. Freiwald

## Abstract

Faces in motion reveal a plethora of information through visual dynamics. Faces can move in complex patterns while transforming facial shape, e.g. during the generation of different emotional expressions. While motion and shape processing have been studied extensively in separate research enterprises, much less is known about their conjunction during biological motion. Here we took advantage of the discovery in brain-imaging studies of an area in the dorsal portion of the macaque monkey superior temporal sulcus (STS), the middle dorsal face area (MD), with selectivity for naturalistic face motion. To gain mechanistic insights into the coding of facial motion, we recorded single-unit activity from MD, testing whether and how MD cells encode face motion. The MD population was highly sensitive to naturalistic facial motion and facial shape. Some MD cells responded only to the conjunction of facial shape and motion, others were selective for facial shape even without movement, and yet others were suppressed by facial motion. We found that this heterogeneous MD population transforms face motion into a higher-dimensional activity space, a representation that would allow for high sensitivity to relevant small-scale movements. Indeed we show that many MD cells carry such sensitivity for eye movements. We further found that MD cells encode motion of head, mouth, and eyes in a separable manner, requiring the use of multiple reference frames. Thus, MD is a bona-fide face-motion area that uses highly heterogeneous cell populations to create codes capturing even complex facial motion trajectories.

## Introduction

The visual world is a dynamic place where many objects are in motion. Some motion patterns, like the visual looming caused by an approaching predator, signifies threats to an animal’s survival (Gibson, 1958). Other motion patterns, like the biological movements of conspecifics can disclose others’ internal states (Puce and Perrett, 2003). Thus, the analysis of biological motion is crucial for agents in a social group. A case in point is facial motion. Faces possess characteristic rigid (e.g. translation or rotation) and non-rigid movements (e.g. dynamic facial expressions). The latter creates both complex visual motion patterns and changes in facial shape. Facial motion is thus challenging for a visual system to analyze. Yet it provides important information, especially for social communication and interaction (Duchaine and Yovel, 2015; Haxby et al., 2000; O’Toole et al., 2002; Pitcher and Ungerleider, 2021).

Where and how is facial motion processed in the brain? Faces are thought to be analyzed by a dedicated face-processing network in the primate brain (Freiwald et al., 2016; Freiwald, 2020; Hesse and Tsao, 2020). This system consists of a set of areas at anatomically stereotypical locations, revealed by functional MRI (fMRI) in both humans and macaque monkeys (Pinsk et al., 2005; Tsao et al., 2003; Tsao et al., 2008). FMRI studies further found that face areas located more dorsally in the superior temporal sulcus (STS), but not face areas located more ventrally, exhibit selective responses to dynamical facial motion (Fisher and Freiwald, 2015; Pitcher et al., 2011). These dorsal areas are thus thought to be specialized for dynamic processing (Duchaine and Yovel, 2015; Pitcher and Ungerleider, 2021).

One such area, the middle dorsal (MD) face area in the macaque brain, showed not only selectivity for facial motion, but for naturalistic face motion (Fisher and Freiwald, 2015): MD responded more to movies of natural face motion than frame-jumbled versions identical on a frame-by-frame basis but with a randomized transition sequence. Ventral face areas like ML and AL showed the opposite pattern of preference (Fisher and Freiwald, 2015). MD’s selectivity for naturalistic facial motion suggests that it may play an important role in facial motion processing. Yet a recent electrophysiological study from MD showed that MD encodes rich facial information purely from static images (Yang and Freiwald, 2021). Some of the static facial information (e.g. facial identity, expression, and gaze) require processing of fine spatial detail. This seems to contrast with the fMRI results as sensitivity to fine spatial detail is uncharacterisitc of motion-processing systems (Nassi and Callaway, 2009). This raises the question whether MD, a hypotetical face-motion area, really does process dynamic facial information at the single-cell level: MD is located next to motion and shape-selecitve areas (Fisher and Freiwald, 2015). It is thus possible that MD does not process facial motion, but might contain two different populations of neurons, one selective to facial shape and the other to general motion. Alternatively, MD could contain cells that are selective for the specific combination of facial shape and facial motion. In this case, MD cells would be integrating facial shape and facial motion, and thus be truly face-motion selective.

Here we aim to determine whether and how MD cells process facial motion. To this end, we localized MD in two macaque monkeys by fMRI, and then recorded single-cell activities inside MD (Yang and Freiwald, 2021). We presented a set of naturalistic and altered face-motion movies to test for face-motion selectivity. We next tested how MD populations encode facial motion and if they were sensitive to physically small, but socially important movements in faces, like gaze shifts. And we were interested in whether MD cells encoded global motion of the head as a rigid body, or if sensitivity to small-scale movements, should it exist, would translate into a more sophisticated representation that was able to separate motion of head, mouth, and eyes.

## Results

### MD cells are selective for naturalistic facial motion

Previous fMRI results (Fisher and Freiwald, 2015) suggest, but do not demonstrate, that MD contains cells selective to facial motion. But such cells have not been found in MD yet. We recorded activity from 178 single units in MD during the presentation of sets of brief 600ms video clips (Figs. 1A and 1B; Move S1; see Methods).

**Fig. 1.**
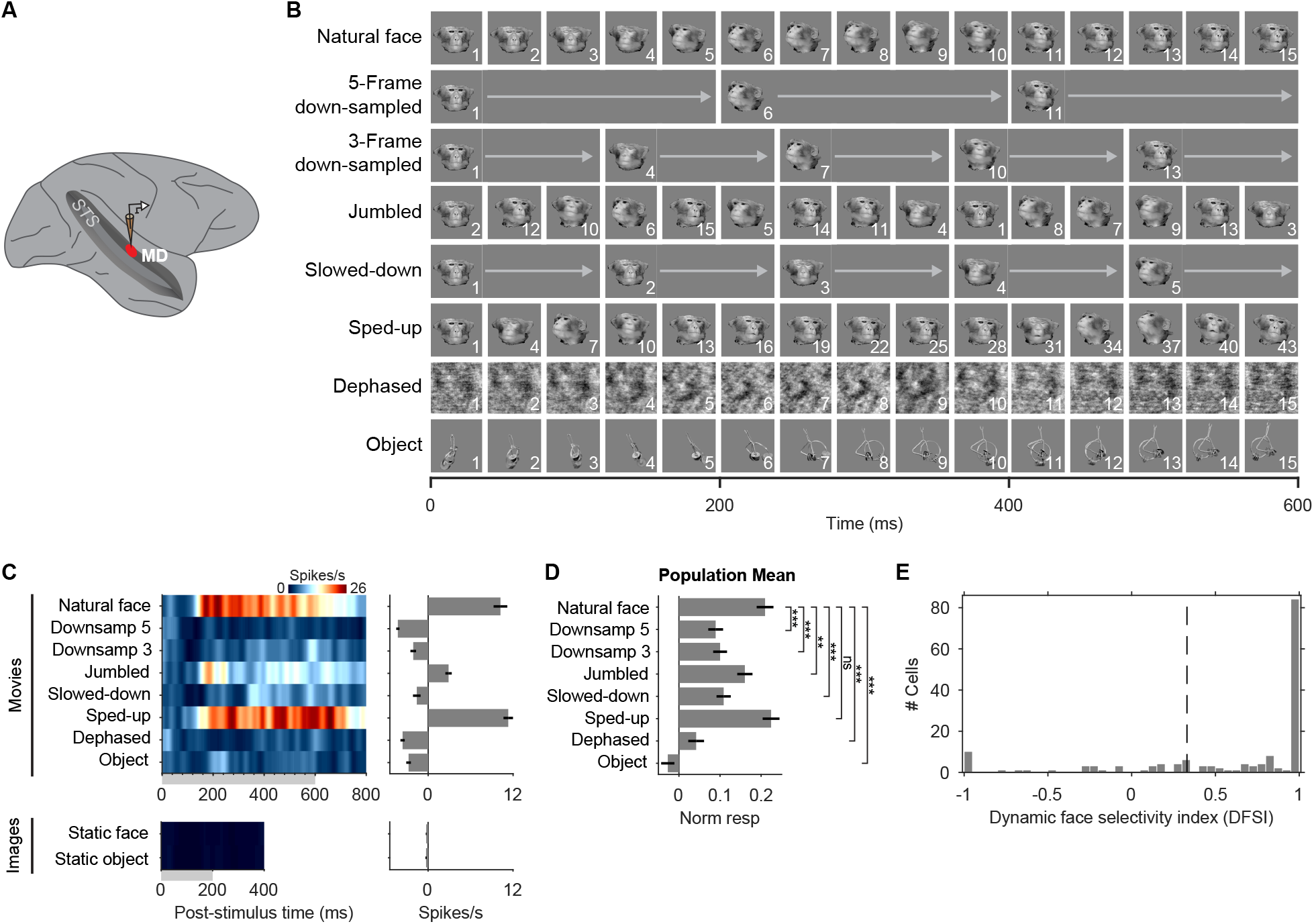
Recordings of MD cells’ responses to dynamic stimuli reveal selectivity to natural face motion. **(A)** Schematic of electrophysiological recording in MD. MD: the middle dorsal face area; STS: superior temporal sulcus. **(B)** Schematic of movies used to test face motion processing in MD. “Natural face” movies were 600-ms videos (25 fps) of monkey facial movements. In the two “down-sampled” conditions, videos were down-sampled to 5 fps and 8.3 fps, respectively. “Jumbled” movies were created by randomizing the frame orders of the natural movies. Slowed-down and sped-up videos were played at speeds three times slower or faster than natural movies, respectively. In “dephased” conditions, we scrambled the phase of the video content by applying the same randomization matrix to the phase component of each frame, which destroyed shape but retained motion. “Object” movies were videos (25 fps) of man-made object movements. Numbers on lower right corner of each image indicate corresponding frames in the natural face condition. See also Movie S1 for example videos. **(C)** Spike density function to movies (top) and static images (bottom) of an example face-motion selective cell. Gray band indicates time periods of stimulus presentation. Minor ticks on x-axis indicate the duration of each movie frame (40 ms). Firing rates (Hz) are color coded. Time averaged response (baseline-subtracted) is shown on the right. **(D)** Mean population responses of all recorded MD cells to dynamic stimuli. \*\**P* < 10^−4^, \*\*\**P* < 10^−10^, ns: *P* > 0.05 (Wilcoxon signed-rank test, false discovery rate corrected). In (C) and (D), error bars represent mean ± SEM. **(E)** Distribution of dynamic face selectivity indices of entire population, calculated from responses to natural face and object movies. Dashed line at 0.33, corresponding to twice as high a response to face than to object movies.

To test facial-motion selectivity for MD cells, we adapted and expanded the paradigm from the previous fMRI study of face-motion selectivity (Fig. 1B) (Fisher and Freiwald, 2015). We included videos of natural face movements, and used down-sampled movies as surrogates for static conditions, while avoiding the effect of neural adaptation to the presentation of long-lasting static images. Comparing these two conditions, we tested for face motion selectivity in MD cells. To test for selectivity for *natural* face motion and to compare with previous fMRI results, we created “jumbled” version of the natural face movies, where the order of frames was randomized, creating an artificial sequence of rapid facial shape changes. We further tested for velocity sensitivity (slowed-down or sped-up movies in Fig. 1B). Finally, we tested sensitivity to shape content by adding “dephased” movies, where frames of natural face movies were phase-scrambled but with low-level motion cues kept unchanged, and object movies.

We found MD cells that behaved similarly as the overall fMRI signal across conditions (Fisher and Freiwald, 2015). In particular, they were strongly selective to facial motion. Figure 1C shows one example. This cell responded vigorously to natural face movies, but was suppressed by the two down-sampled counterparts (Fig. 1C, top). Yet this cell did not simply respond to any dynamic stimulus - it was highly selective to motion content. Its response was reduced to less than one third of that to natural face movies, when presented with “jumbled” face movies. The cell was also sensitive to the velocity of facial motion, with suppressed responses to slowed-down movies, and slightly increased responses to sped-up movies. Finally, the motion selectivity of this cell was specific to faces, as it was suppressed by “dephased” and object movies. In total, these results indicate that the cell requires the presence of both facial shape and natural facial motion to respond, thus is truly face-motion selective.

As a further test of static shape selectivity, for this and all other cells recorded this way, we compared their responses to static images as measured in an independent experiment (see Methods). In brief we presented static images of faces and non-face objects for 200 ms followed by 200 ms of blank screen. The example cell in Fig. 1C did not respond at all to static faces or objects (Fig. 1C, bottom). Thus this cell only responded to dynamic movies, but not static images.

Across all recorded cells, the MD population showed a similar response pattern as the example cell. On average, the MD population responded about twice as strongly to natural face movies than to both down-sampled and slowed-down movies (Fig. 1D; normalized responses: 0.21 for natural face movies; 0.09, 0.10, 0.11 for 5-frame down-sampled, 3-frame down-sampled, and slowed-down movies respectively), suggesting that MD neurons are sensitive to naturalistic face-motion velocities, but do not encode velocity per se. Supporting this view, MD neurons responded to faster movies at about the same magnitude (normalized responses: 0.22 for sped-up movies) as to the original naturalistic movies. MD neurons were highly sensitive to the natural sequence of the movie frames as opposed to the shape content alone, responding significantly more to naturalistic movies than to jumbled ones (normalized responses: 0.16 for jumbled movies; *P* <10^−4^, Wilcoxon signed-rank test, false discovery rate [FDR] corrected). Finally, MD neurons were also sensitive to the shape content of the movies, and not just its motion content, responding minimally or even being suppressed during presentation of the two motion conditions devoid of facial shape (“dephased” and “object”). Thus, the MD population as a whole is both facial shape and motion selective.

We next asked how face selective MD neurons were in dynamic settings. In our previous recordings in MD (Yang and Freiwald, 2021), we had found that MD cells have high face selectivity to static images. Does this selectivity extend to dynamic settings? We quantified the degree of dynamic face selectivity of all recorded cells with a dynamic face selectivity index (DFSI) in analogy to the static face selectivity index. The DFSI compares responses to natural face movies *R*_*natural*_ with those to nonface object movies *R*_*object*_: *DFSI = (R*_*natural*_ *– R*_*object*_*) / (R*_*natural*_ *+ R*_*object*_) (see Methods). The MD population is highly face selective for dynamic stimuli, with the distribution of the DFSIs heavily skewed to the right (Fig. 1E; mean DFSI=0.61). In total, 76% cells were dynamic face selective, responding at least twice as strongly to face motion than to object motion. These numbers are similar to those during the presentation of static images (84% of MD cells responsive to static images are static face selective). Thus the MD population exhibits similarly high degrees of facial shape selectivity in static and dynamic conditions.

### MD exhibits a high heterogeneity of tuning to naturalistic facial motion

The mean MD population response exhibited a clear-cut preference for naturalistic (and sped-up) face-containing movies. While many cells, like the example in Fig. 1C, showed a tuning profile similar to that of the population (Fig. 1D), other cells exhibited a very different tuning to facial motion. Figure 2A shows an example cell with an almost complementary response profile. This cell responded strongly to down-sampled or slowed-down face motion, but barely to natural, jumbled, or sped-up face motion (Fig. 2A, top). Its initial responses to these three conditions were to the first movie frame, and once the second frame started creating facial motion (40 ms after onset of the first frame), the response was almost completely abolished. The cell’s response to static images (Fig. 2A, bottom) shows that it was static face selective, which is independently confirmed by its weak or suppressed response to the two nonface motion conditions. Thus this neuron is face-shape selective and suppressed by facial motion.

**Fig. 2.**
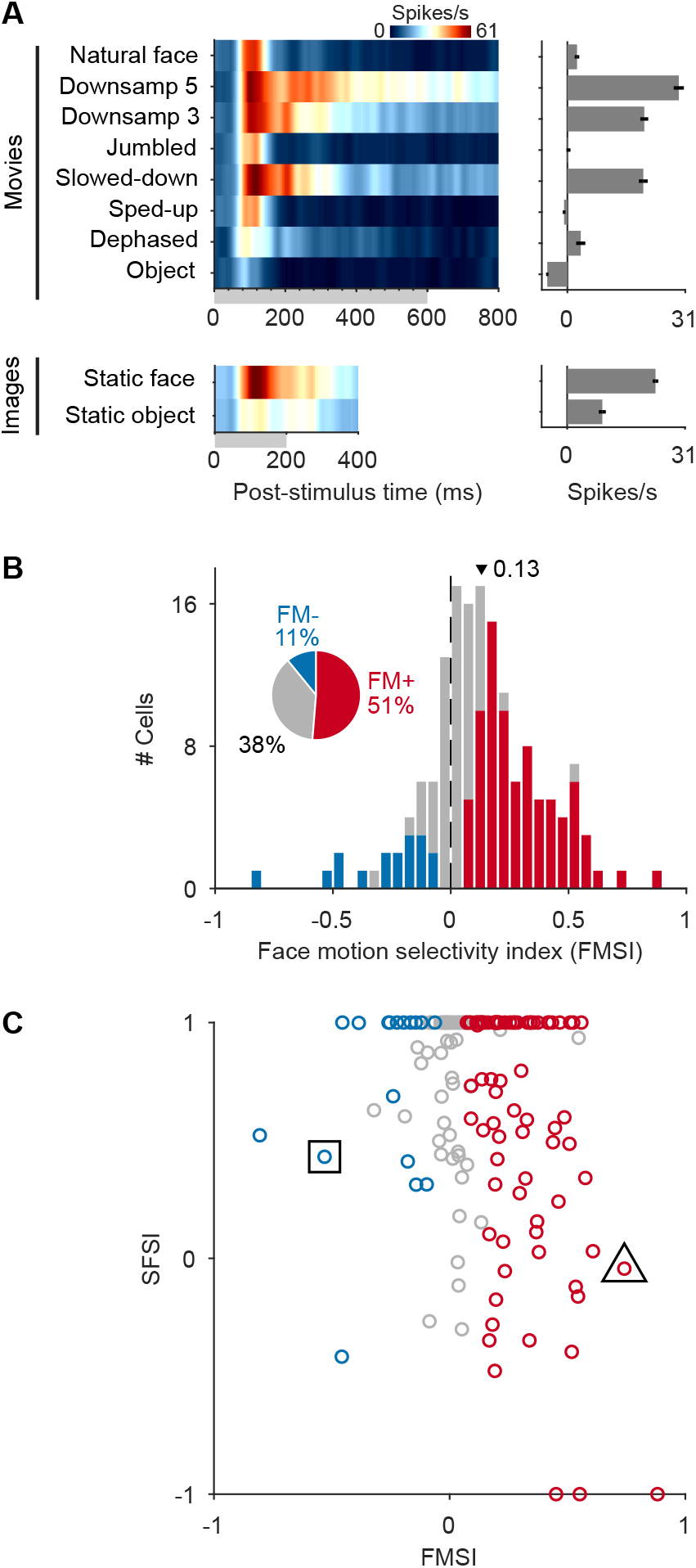
MD shows highly heterogenous responses to facial motion. **(A)** Spike density function to movies (top) and static images (bottom) of an example face-motion suppressive cell. Gray band indicates time periods of stimulus presentation. Minor ticks on x-axis indicate the duration of each movie frame (40 ms). Firing rates (Hz) are color coded. Time averaged response (baseline-subtracted) is show on the right. Error bars denote mean ± SEM. **(B)** Distribution of face motion selectivity indices (FMSIs) across all face-selective cells. Proportion of cell tuning was inserted as the pie chart. Top triangle depicts mean FMSI. FM+: face motion preferred, FM-: face motion suppression. Red and blue denotes FM+ cells FM-cells respectively, and gray denotes cells that are not tuned. **(C)** Scatter plot of static face selectivity index versus FMSI for all face-selective cells. Triangle and square mark example cells in Fig. 1C and Fig. 2A respectively. Color coding as in (B).

To quantify the range of preferences for facial motion, we calculated a facial-motion selectivity index (FMSI) for each face-selective cell (including both dynamic and static face selective ones; 156 neurons in total), by comparing its response to natural face motion (*R*_*natural*_) with that to the 5-frame down-sampled (*R*_*downsamp5*_) version: *FMSI = (R*_*natural*_ *-R*_*downsamp5*_*) / (R*_*natural*_ *+ R*_*downsamp5*_) (see Methods). The distribution of FMSIs shows that many MD neurons show a significant preference for facial motion (Fig. 2B; population mean ± SD: 0.13 ± 0.24). Statistical tests (Wilcoxon rank-sum tests; *P* < 0.05; see Methods) revealed that that 80 (51%) of face-selective MD cells responded significantly more strongly to faces in motion (FM+ cells; Fig. 2B, insert), while 17 (11%) were selectively suppressed by facial motion (FM-cells; Fig. 2B, insert), and 59 (38%) appeared to be insensitive to facial motion. Thus static and moving faces will engage different MD populations, and a moving face stimulus will not only increase, but also diversify the MD population response.

The example cells in Figs. 1C and 2A might suggest that there could be a strict separation between cells that are static face selective and those that are face motion selective. We tested this by plotting each cell’s static face selectivity index (SFSI, see Methods) against its FMSI (Fig. 2C). We found that selectivity for facial shape and face motion were largely independent across cells (Fig. 2C; Spearman’s *ρ* = -0.17, *P* = 0.03, permutation test). This indicates that MD cells could show face-motion selectivity no matter whether they are highly selective for static facial shape or not. Similarly, static face-selective cells could also show (or not show) face-motion selectivity. Thus, there is great heterogeneity of face and motion coding in MD, but it is not of the kind predicted by a two-population model of MD with one sub-population generically motion-, but not face-selective, and the other one facial-shape but not motion selective. This model would predict low absolute SFSI and high FMSI values for population one, and high SFSI but low FMSI values for population two. Figure 2C does not show such two separated clusters with these combination of values. The model would also predict a strong negative correlation between SFSI and FMSI across the two populations, which is neither apparent in MD (Fig. 2C).

### The MD population response captures physical motion similarities and represents face motion in a higher-dimensional space

How does MD population encode facial motion? To address this general question, we first asked whether the MD population response is sensitive to the optic flow in dynamic faces. We estimated optic flow between natural face movie frames for all pixels (Fig. 3A). We then projected, via a principal component (PC) analysis, the optic-flow data into a low-dimensional space (Fig. 3B, bottom). We then applied the same dimensionality reduction to the population responses of all FM+ cells (Fig. 3B, top). We found that physical and neural inter-movie distances were highly correlated across movies (Fig. 3C; Pearson’s *r* = 0.62, *P* <10^−7^). This result shows that FM+ neurons in MD capture the main physical motion dynamics. We performed the same analysis for object motion videos, but did not find any correlation of optic flow and the FM+ population response (Fig. 3D; Pearson’s *r* = 0.08, *P* = 0.51). Thus the FM+ population response is specific to motion patterns of faces, not motion patterns in general.

**Fig. 3.**
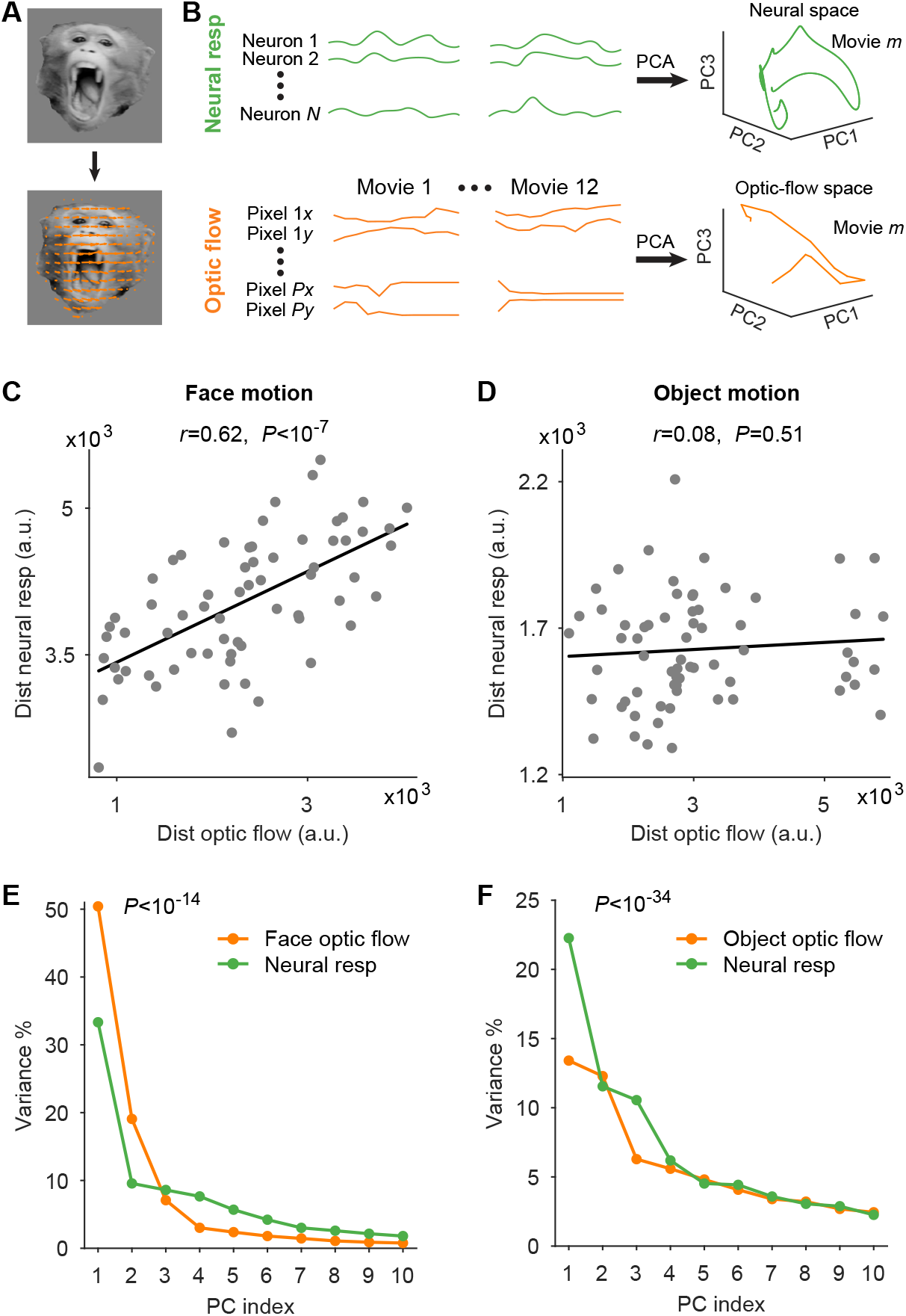
MD captures physical similarities of face motion and encodes facial dynamics by dimensionality expansion. **(A)** Example movie frame (top) and the estimated optic flow (orange arrows) overlaid on the frame (bottom). **(B)** Schematic of dimensionality reduction analyses on optic flow and neural responses. First three principal components (PCs) are shown for demonstration. **(C)** For face motion, distances between movie trajectories in optic-flow space were correlated with distances in the neural space (*r*: Pearson correlation). **(D)** For object motion, distances between movie trajectories in optic-flow space were not correlated with distances in the neural space. **(E** and **F)** Distribution of variance explained by first ten PCs in optic-flow and neural spaces for face motion (E) and object motion (F). Broader distribution suggests higher dimensionality. In (E) and (F), *P* values are from Kolmogorov-Smirnov tests.

The PC analyses also allowed us to characterize the dimensionality of both. If the FM+ population only captured the main motion patterns, it should have the same dimensionality as the flow pattern itself. If it would not capture all of the dynamics, it should be lower-dimensional, and if it was sensitive to finer stimulus details which are coded by lower-rank PCs (Stringer et al., 2019), it should be higher-dimensional. The comparison of the PC power spectra for optic flow and population response revealed that the population response was higher-dimensional than the physical optic flow (Fig. 3E; *P* <10^−14^, Kolmogorov-Smirnov test): while 69% variance of the optic flow patterns was explained by the first two PCs, they explained only 43% variance of the neural population response. Furthermore, the neural PC spectrum remained flatter for higher PCs than that of the optic flow (Fig. 3E). As a result, it requires more than twice the number of PCs to explain at least 80% of total variance for neural data (11 PCs) compared to optic flow (5 PCs). We did not find this relationship in the case of object motion (Fig. 3F). Here the dimensionality of the population response was slightly smaller than that of the optic flow pattern (*P* <10^−34^, Kolmogorov-Smirnov test). Thus the FM+ population expands the dimensionality of representation specifically during the encoding of face motion stimuli. How could this dimensionality expansion be used? One possibility is that physically less prominent motion is emphasized.

### MD exhibits sensitivity to small movements of the eyes

In faces, one physically small but socially important motion signal is the shift of gaze direction, which signifies the reorientation of attention of another subject (Carlin and Calder, 2013). We previously found that MD contains cells tuned to static gaze direction, indicating MD cells are sensitive to small differences shown in the eye region (Yang and Freiwald, 2021). This raises the possibility that MD cells might also signal dynamic gaze shifts, even though these shifts are physically far less prominent than the motion of the head or the mouth. To this end, in a second experiment we created minimally dynamic stimuli in which the face was kept still except that the gaze switched from one direction to another (Fig. 4A; see Methods; Movie S2). We created these stimuli for three different head orientations (Fig. 4A).

**Fig. 4.**
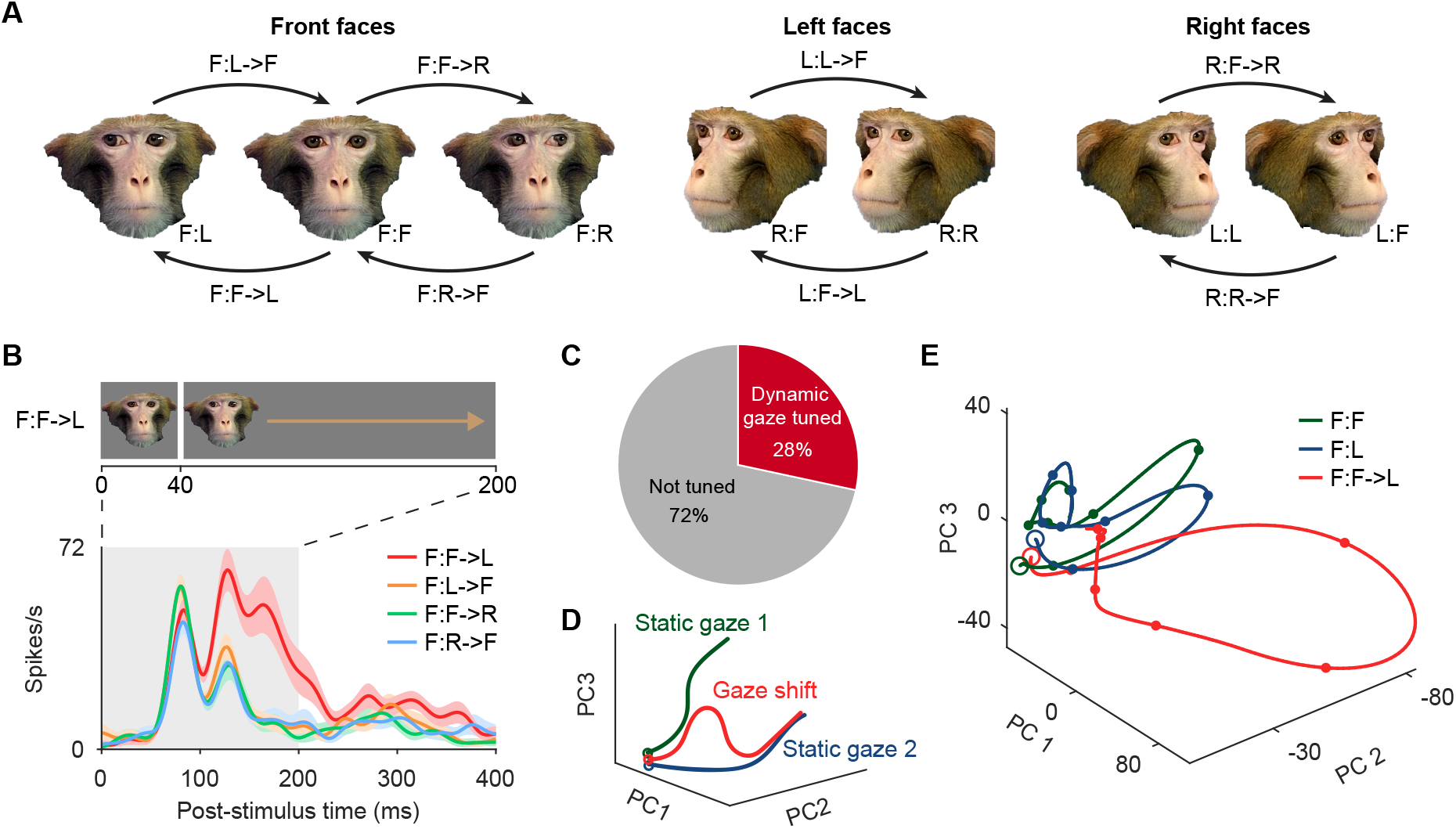
MD is sensitive to small movement of the eyes. **(A)** Example stimuli of dynamic and static gaze conditions with condition labels. Dynamic stimuli were gaze shifts (indicated by arrows) between two static gaze conditions. Dynamic conditions are labeled as “head orientation : first gaze direction -> second gaze direction”. Static conditions are labeled as “head orientation : gaze direction”. L: left, F: front, R: right. **(B)** Spike density function of an example cell showing tuning to gaze shifts. Shading of each line denotes mean ± SEM. Gray band illustrates time period of stimulus presentation. (Top) Temporal structure of an example gaze shift: one frame (40 ms) of first static gaze followed by 160 ms of second static gaze. **(C)** Distribution of face-selective cells tuned to gaze shifts. **(D)** Hypothesized neural dynamics to gaze shifts: a gaze shift could induce a transition of state from the neural trajectory to the first static gaze to the trajectory to the second static gaze. Open circles denote starting states. **(E)** Neural population trajectories of gaze-tuned cells to one example gaze shift condition (red) and the two composing static gaze conditions shown in neural space of first three principal components. Open circles denote starting states and filled dots indicate 40-ms intervals. (See Fig. S2 for results of all other gaze shift conditions).

Figure 4B shows an example cell with tuning to gaze shifts. When presented with front-view faces (“F:” in figure legend), gaze direction could change in four different ways, from front to left (F->L), from left to front (L->F), from front to right (F->R), and from right to front (R->F). The cell responded to three out of these four conditions virtually indistinguishably, but significantly differently to one: a shift of the eye position from front to left elicited a much bigger and longer-lasting response. Note that this means that the cell did not just respond to a particular movement direction, since two conditions (F:F->L and F:R->F) contained the same motion direction, but the cell responded to only one. On the other hand, the cell’s tuning to dynamic gaze shifts could not be explained by tuning to static gaze directions, since its tuning became prominent only in the second response phase after the gaze had switched. Thus, this kind of selectivity can only be explained by a conjunction of detailed shape and motion analysis.

Comparing the responses to different gaze shifts, we found that out of the 141 face-selective cells recorded in this experiment, 40 (28%) showed significant modulation by gaze shifts within at least one head orientation (Wilcoxon rank-sum tests for half profile viewpoints or Kruskal-Wallis tests for front viewpoints; *P*<0.05, FDR corrected). Among these cells, 27 (68%) of the 40 gaze-shift tuned cells were not tuned to static gaze at all (Fig. S1). Thus their tuning to gaze shifts is not a byproduct from static gaze tuning, indicating that specific subpopulations of MD cells encode static and dynamic gaze.

We next asked how the MD population captures such gaze shifts. The simplest possibility is an update model: different gaze directions are encoded by different population response trajectories, and a gaze shift by the transition from one trajectory to the other: the population response is updated from the beginning to the ending trajectory (Fig. 4D). Alternatively, the population could generate an entirely new activity trajectory, highlighting the occurrence of a change in gaze direction and differentiating it from conditions without gaze change.

To test this, we studied the neural dynamics by performing a PC analysis on the populational responses to all gaze stimuli, including dynamic and static gaze. We then visualized the neural dynamics by projecting population responses to each gaze shift and the two composing static gaze directions onto the same first three PCs. Figure 4E shows the neural trajectories to the gaze shift from front to left within a frontal face (red curve), and to static front (green curve) and to static left (blue curve) gaze directions, also in a frontal face. The population response to the gaze shift, it turned out, was not a simple transition between the two static responses, but rather bifurcated from the static-gaze responses into a new state space (Fig. 4E). We found this pattern of trajectories for dynamic stimuli separating from the trajectories of component static conditions for all eight gaze shifts we studied (Fig. S2). The data thus refute the updating model. Rather, MD, with exquisite sensitivity to the physically small changes in the face induced by gaze changes, represents gaze shifts in a qualitatively different way from static gaze.

### MD represents the motion of multiple face parts, using multiple reference frames

We have found that the MD population represents very different scales of motion from the optic flow patterns of the entire moving face to the small changes of the eyes within a static face. The anatomy of the primate face allows for independent movements of the head and its parts, in particular the eyes and the mouth. Consider, for example, a threat gesture in which the head is turned towards an opponent, while the mouth is opened. We thus wondered whether MD cells would be able to capture these different forms of facial movement and, if so, how.

For a computational system to be able to represent these different movements and their combinations and thus to fully represent face motion, it needs to de-mix the motion of facial parts from global head motion. In other words, the system would need to create different reference frames: a viewer-referenced frame for global motion of the head, and face-referenced frames for part-based motion of the eyes and the mouth. Furthermore, the system would need to be able to represent these different movements despite their vastly differing amounts of motion energy (head > mouth > eyes).

We tested whether MD cells can accomplish this nuanced representation of facial motion. To this end, we first tracked facial landmarks in the natural face movies using machine learning (Fig. 5A; see Methods). Based on the tracking result, we created regions of interest (ROIs) for different facial motion components. This allowed us to estimate the distribution of the motion energy of head, eye, and mouth motion separately (Fig. 5A and 5B; see Methods; Movie S3). As expected, global motion of the head was estimated to have the highest energy, followed by motion of the mouth, while motion of the eyes was the weakest (Fig. 5B).

**Fig. 5.**
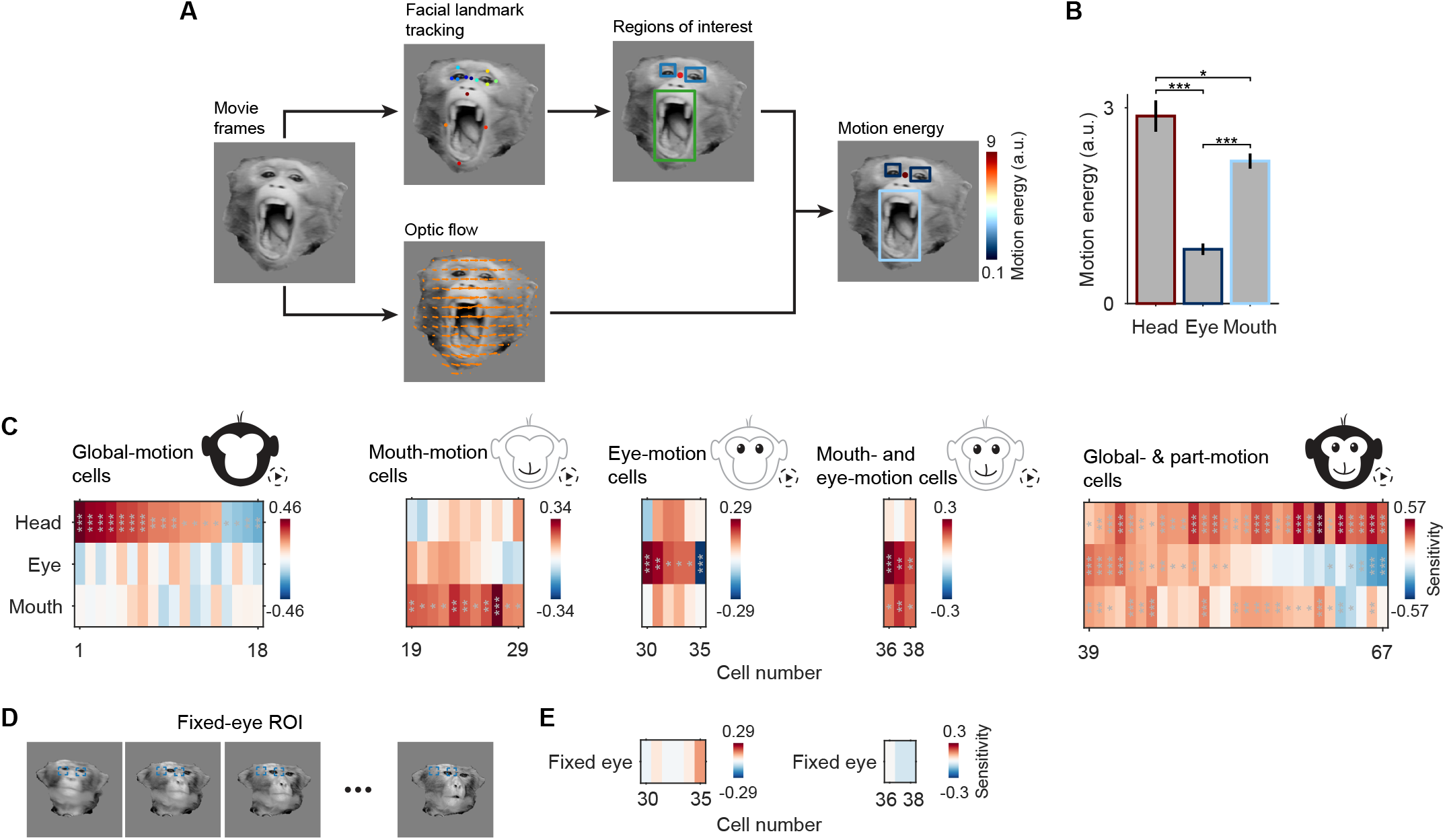
MD encodes face motion through multiple reference frames. **(A)** Workflow for motion energy estimation. For each movie, facial landmarks were tracked using DeepLabCut (Mathis *et al*., 2018; Nath *et al*., 2019). Regions of interest (ROIs) were then created based on the tracked landmarks to capture global head motion, motion of the eyes and of the mouth. Motion energy of the head was estimated by the magnitude of the averaged optic-flow vector (bottom) within the head ROI. Within the eye and mouth ROIs, the head optic-flow vector was first subtracted to get the head-referenced motion vectors, after which the vector magnitude was averaged to obtain the energy values for the eyes and mouth. See Movie S3 for full results of motion estimation. **(B)** Averaged motion energy of the head, eyes, and mouth across natural face movies. Error bars indicate mean ± SEM. \**P* < 0.05, \*\*\**P* < 10^−22^ (Wilcoxon signed-rank test, false discovery rate corrected). **(C)** Face-motion sensitivity matrices for cells showing sensitivity to global head motion (leftmost panel), to face part motion of the eyes or mouth (middle three panels), and to motion of both head and eyes or mouth (rightmost panel). Color coded is the neural sensitivity – standardized regression coefficients – of each cell to each motion energy. Overlaid stars mark significant sensitivity. \**P* < 0.05, \*\**P* < 0.01, \*\*\**P* < 0.001 (*T*-test). **(D)** Schematic of the control analysis for eye motion sensitivity. The fixed-eye ROIs were fixed rectangles at the averaged position of the original eye ROIs. **(E)** Sensitivity to fixed-eye motion energy for eye-motion sensitive cells in (C). Conventions as in (C). No eye-motion sensitive cells showed significant sensitivity to fixed-eye motion.

We then modeled the spike train of each FM+ neuron by linear regression of the three motion components (see Methods) and quantified the sensitivity of each cell to these three components of facial motion by the regression coefficients (Fig. 5C). We found that 23% (n=18; Fig. 5C, leftmost panel) of cells were sensitive to motion of the head alone, 25% (n=20) to either motion of the mouth or eyes, but not to that of the head (Fig. 5C, three middle panels). About a third of the neurons (36%; n=29) were sensitive to global (head) and different combinations of part (eye, mouth) motion simultaneously (Fig. 5C, rightmost panel). Thus MD cells have the capacity to encode global head motion and to encode local motion of face parts while ignoring global head motion. The latter requires the tracking of facial parts in a reference frame moving with the head.

To test whether part-motion sensitive cells were truly tuned to local motion, we performed two control analyses. We focused on the eye region given that its small signal is particularly susceptible to contamination from other sources of facial motion. First, we tested whether the eye motion signal we estimated represents purely the local motion of the eyes or might be contaminated by components of global head motion. To this end, we created a shift-control in which we moved the eye ROI to a position just outside of the eyes, while otherwise, except for this offset, preserving the precise motion path of the original ROI (see Methods). This shift-ROI would thus estimate not local eye-motion information, but rather a potential nearby global-motion contamination source. If this motion component did not contribute to our original estimation of local motion, the new ROI should estimate motion energy close to zero. This is exactly what we found (Fig. S3). The original eye ROI already had very weak motion energy (Fig 5B), but for the new ROI, we found a further 55% reduction in motion energy compared to the original ROI (mean motion energy: 0.83 for the original eye ROI and 0.38 for the shifted ROI). Second, we tested the idea that a head-centered reference frame is necessary for eye-motion sensitive cells to be activated. In this control analysis, we placed the eye ROIs at a fixed position (Fig. 5D; see Methods) and re-estimated the motion energy within these “fixed eye” ROIs. We found that there was a lot of stimulus motion energy in this new ROI, much more than to the eyes in the original computation (Fig. S4). Yet critically, none of the eye-motion sensitive cells were sensitive to this motion energy (Fig. 5E). Thus these cells are specifically sensitive to eye motion within a face-centered reference frame.

In sum, these results imply that the population of MD cells can represent motion of eyes, mouth, and head separately, and that they do so by establishing separate reference frames.

## Discussion

Faces constitute a special class of visual objects with a shared three-dimensional structure. Faces, like other biological shapes, but unlike most inanimate ones, can change their shape dynamically, creating complex shape-motion dynamics. This physical duality of faces implies a duality in the kinds of information faces can convey to visual systems that are equipped to extract both the immutable, structural properties revealing qualities like facial identity, as well as the changeable and changing properties revealing qualities like facial expressions.

Since the now-classical functional face-processing model by Bruce and Young (Bruce and Young, 1986), it is widely assumed that these two dimensions of faces are processed by different systems. Following Bruce and Young (1986), the Haxby model (Haxby *et al*., 2000), proposed a ventral processing pathway to extract structural properties of faces, and a dorsal one for processing changeable or changing dimensions of faces. These models have stood the test of time and been confirmed and extended by innumerous imaging studies (Duchaine and Yovel, 2015; Fisher and Freiwald, 2015; Pitcher *et al*., 2011; Pitcher and Ungerleider, 2021; Polosecki et al., 2013). But how are the different qualities of faces extracted from incoming visual stimuli?

Much of the knowledge we have obtained to answer this question, comes from experiments combining whole-brain fMRI to localize face-selective areas in the macaque monkeys with targeted electrophysiological recordings of single cells from these areas (Tsao et al., 2006). This line of work has focused on the extraction of facial identity information in the face of variations of size, position, and head orientation (Chang and Tsao, 2017; Freiwald and Tsao, 2010; Freiwald et al., 2009; Koyano et al., 2020; Tsao *et al*., 2006). The work describes a transformation of representations from an initial strongly head-orientation-selective one to a strongly identity-selective one along multiple face areas. Similar mechanistic insights are lacking for the second major source of facial information, facial dynamics.

The discovery of dorsal face area MD and its functional signature of facial motion selectivity (Fisher and Freiwald, 2015), occurred much later than the discovery of ventral face areas (Pinsk *et al*., 2005; Tsao *et al*., 2003; Tsao *et al*., 2008). It opened the possibility to use the same approach of fMRI-guided single-cell electrophysiology to gain similar mechanistic insights into the encoding of facial dynamics. Our study is the first to do so. There are only two prior single-unit studies on tuning to face and body movements (Oram and Perrett, 1996; Perrett et al., 1985), which focused on rigid body and face movements, and these studies were not fMRI-guided. Based on the anatomical information provided in these papers, many of the recordings were taken anteriorly to the ones in this study.

MD is located in the dorsal bank of STS and is thought to be the first area of the dorsal stream of the face-processing network, homologous to the human posterior STS face area (Fisher and Freiwald, 2015; Weiner and Grill-Spector, 2015; Yang and Freiwald, 2021). Previous fMRI studies in humans and monkeys have found that dorsal face areas show much stronger responses to dynamic faces compared to static ones (Fisher and Freiwald, 2015; Pitcher *et al*., 2011; Polosecki *et al*., 2013). MD has been further shown in fMRI experiments to be selective to natural face motion, and not generally to changes of the face, e.g. jumbled face motion (Fisher and Freiwald, 2015). This made MD the most promising target for a single-cell investigation of the processing of facial dynamics. Here we made four major discoveries.

First, because MD is located directly neighboring generally motion-selective areas (Fisher and Freiwald, 2015), and because fMRI has limited spatial resolution, fMRI results could not discern between genuine processing of facial dynamics and a two-population model of separate face and generic motion cells. According to the latter scenario, MD would not really constitute a face-motion area. Here we show that it is. Four main observations support this view: (a) the mixing model predicts sizeable responses to non-face motion, yet we did not find those (Fig. 1D). (b) the mixing model predicts a large population of non-face selective neurons during presentation of dynamic stimuli, yet we found the majority of MD cells maintained face-selective under these conditions (Fig. 1E). (c) the mixing model predicts two clusters of cells with separate combinations of FMSI and SFSI, which we did not find (Fig. 2C). (d), we found cells that responded to dynamic faces, and neither to static faces nor non-face motion (Fig. 1C). This is the most direct evidence that MD truly integrates facial shape and motion.

Second, our study provides first insights into the nature of the population code for face motion. We found that MD shifts the representation of facial dynamics into a space that is higher dimensional than physical motion space. Expansions of dimensionality have been seen in the visual system before, most dramatically in the more than 100-fold increase in numbers of neurons in striate cortex compared to the lateral geniculate nucleus (LGN) (Connolly and Van Essen, 1984). Here we show such dimension expansion in the visual processing of complex object motion. We hypothesized that the increase in dimensionality could result from MD emphasizing physically small but behaviorally important facial motion and indeed found MD cells that are highly sensitive to the physically small movements of the eyes. Dimensionality expansion could yield further benefits. It might help MD disentangle the highly nonlinear face motion pattern into a more separable representation, similar to the principles behind kernel-based support vector machines, which project non-linear data into higher dimensional space to make them linearly separable (Boser et al., 1992).

Third, we discovered that MD cells are sensitive to different components of facial motion. Many MD cells are sensitive to the movements of the head. But there are also MD cells that are sensitive to the local motion of the eyes or the mouth, while ignoring the global motion of the head. This suggests that these cells build a representation of the movements of these facial parts *relative* to the head in the movies, i.e. within an object-referenced coordinate system. This is in contrast to head-motion sensitive cells, which only requires a viewer-referenced frame. Thus MD might contains two different reference frames for face motion, and may perform a transformation of coordinates when representing the motion of facial parts.

What computations might underly this ability? Now-classical work on multiple reference frames in parietal cortex (Snyder et al., 1998), described gain fields. Gain modulation integrates two sources of information with one input changing the gain of the neural response to the other input (Salinas and Abbott, 2001; Salinas and Sejnowski, 2001; Zipser and Andersen, 1988). Such gain modulation has been proposed to serve reference frame transformations rather generally in sensory-motor transformation and navigation (Salinas and Thier, 2000). We propose it as a potential mechanism underlying the sensitivity to local motion of facial parts independent of head motion direction. MD cells sensitive to head orientation (Yang and Freiwald, 2021) could provide the gain fields needed for this operation.

Fourth, MD contains a high heterogeneity of selectivity to face motion: while about half of the face cells in MD preferred face motion, the others do not prefer face motion or are even suppressed by the presence of motion; some cells are selective for head motion, others for the mouth, and yet others for the eye, and many encode combinations. Further adding to this complexity is the heterogeneity of tuning to facial shape, which we described in a previous study (Yang and Freiwald, 2021) in the same population of MD cells. MD cells exhibit a wide range of tuning to head orientation, gaze direction and the combination of head and gaze direction, and they are also selective for facial expression and facial identity, also in varying combinations. Such diverse tuning to so many dimensions of facial properties has not been described in other face areas so far. In particular, selectivity to facial expression, gaze, and facial dynamics, to the best of our knowledge, have not been studied in ventral face areas yet (Chang and Tsao, 2017; Freiwald and Tsao, 2010; Freiwald *et al*., 2009; Koyano *et al*., 2020; Tsao *et al*., 2006). And in ventral face areas tuning to head orientation progresses in an orderly manner from one face area to the next rather than, as in MD, mixing within a single face area (Freiwald and Tsao, 2010).

Thus our results pose a major challenge to current thinking about the computations performed by face-processing circuits. Computational models of face-processing have focused on the hierarchical nature of one major transformation, which shallow and deep neural networks can perform, resembling the neural population codes at three levels of the face-processing hierarchy (Leibo et al., 2017; Yildirim et al., 2020). Computational models of MD, in turn, cannot focus on a single operation, and must grapple with the fact that multiple computations could have rather different requirements. We alluded to the possibility of gain field like operations to compute multiple reference frames. This needs to operate on top of computations integrating shape and motion, which might occur via AND-like operations on static face-selective and facial motion inputs (Giese and Poggio, 2003; London and Hausser, 2005), all the while multiple dimensions of faces are extracted as well. These are formidable challenges for a computational understanding of MD.

What we do understand though is that these results challenge the major theoretical account of face processing systems with a ventral one focusing on structural properties of faces and a dorsal one focusing on changeable or changing aspects of faces. First, and most importantly, the high heterogeneity of tuning to structural, changeable, and changing properties of faces in a dorsal face area, challenges the fundamental dissociation these theories propose for the two pathways. Even without knowing the selectivity of ventral areas to changeable dimensions of faces like gaze direction or facial expression and to facial dynamics, it is clear that the functional dissociation between pathways is different to the one originally proposed – or more quantitative than qualitative in nature (Bruce and Young, 1986; Calder and Young, 2005; Duchaine and Yovel, 2015; Haxby *et al*., 2000; Pitcher and Ungerleider, 2021). Second, the debate about whether the dorsal stream encodes changeable or changing features of faces (Duchaine and Yovel, 2015), is resolved, at least for the macaque monkey brain. We show that MD is exquisitely tuned to both of these dimensions of faces.

Thus, this study lays the foundation for the physiological understanding of the processing of facial dynamics. Just as ventral face areas have served as a model system for object recognition in general (Freiwald, 2020; Hesse and Tsao, 2020), area MD may serve as a model system to gain mechanistic insights for the field of shape-motion processing, which has received much less attention in the past (Fisher and Freiwald, 2015). Our findings require new thinking about computational models underlying face processing in general and new functional accounts extending the currently dominating ones.

## Materials and Methods

Extra-cellular electrophysiological data were recorded from two male rhesus monkeys (*Macaca mulatta*) between 8 and 10 years old. We first localized MD using fMRI and then guided our recordings by fMRI activations. During each recording session, a tungsten electrode (FHC) was back-loaded into a metal guide tube, and then advanced slowly by hand using a Narishige microdrive. After the electrode reached the desired depth according to high-resolution MRI scans, well-isolated units were recorded from MD. Neural data were sampled at 30 kHz and recorded by a Blackrock acquisition system (Blackrock Microsystems). Action potentials were extracted online using a set of amplitude hoops and were re-sorted offline using the BOSS software (Blackrock Microsystems). See also Yang and Freiwald, 2021 (Yang and Freiwald, 2021) for details about surgeries, MRI and fMRI localization, and electrophysiological recordings.

All procedures followed the *NIH Guide for Care and Use of Laboratory Animals*, and all experiments were performed under the approval of the Institutional Animal Care and Use Committees (IACUC) of the Rockefeller University.

### Behavioral Tasks and Visual Stimuli

Head-fixed subjects were kept in a Faraday cage and fixated on a small dot on a CRT monitor at a distance of 57 cm. The fixation dot had a size of 0.16 degree of visual angle (dva), and fixation window was 2.5 dva × 2.5 dva. The monitor covered a visual field of 39.2 dva × 29.9 dva. Stimuli were presented at a 100 Hz refresh rate in random order. An ISCAN system tracked eye position at 120 Hz. The timing of stimulus presentation was recorded by a photodiode placed on the lower right corner of the screen.

#### Dynamic faces

The stimulus set consisted of 96 grayscale videos clips (600ms) in eight categories: “natural face”, “5-frame down-sampled”, “3-frame down-sampled”, “jumbled”, “slowed-down”, “sped-up”, “dephased”, and “object” (Fig. 1B, Movie S1). The “natural face” condition contained 12 videos (25 fps) of monkey facial movements. These movies were shot with monkeys sitting in vertical chairs. Backgrounds and chair surfaces were covered with blue cardboards or tape, which were removed during post-processing by the chroma key tool in Final Cut Pro (Apple). Stimuli were rescaled to the same size, and luminance across movie frames was histogram-normalized. The “object” condition was composed of 12 videos of moving objects that the subjects were familiar with. Stimuli in all other categories were derived from the natural face movies using MATLAB (Mathworks). Videos in the two down-sampled categories were made by down-sampling the natural face videos to 5 fps (5-frame down-sampled) and 8.3 fps (3-frame down-sampled), respectively. Jumbled videos were created by randomizing the order of frames. The sped-up videos were generated by skipping every second and third frames, while the slowed-down videos were created by repeating each frame three times. Dephased videos were produced by first calculating the Fourier transformation of the frames, and then applying a random transformation matrix to the phase component of each frame. The same random transformation matrix was applied to the frames of the same video, and the order of frames was kept, so that luminance and motion energy remained unchanged in the dephased compared to original videos. Movies (14 dva × 14 dva) were presented at the center of the screen and were separated by 200 ms of blank intervals.

#### Static faces and non-face objects

We also recorded neural responses to a stimulus set of static images including faces, bodies, and non-face objects (the FOB stimulus set) to compare static and dynamic processing in MD from the same cells in a previous study (Yang and Freiwald, 2021). Description of the stimulus set, as well as corresponding neural responses, have been described in detail before (Yang and Freiwald, 2021). Briefly, the stimulus set contains six categories of static images, including monkey and human faces, monkey and human bodies, fruits, and manmade objects. Stimuli (14 dva × 14 dva) were presented for 200 ms, followed by 200 ms of gray screen.

#### Dynamic and static gaze

Dynamic gaze conditions consisted of 24 monkey face videos (200 ms, 25 fps) from three identities with three head orientations (Fig. 4A, Movie S2). Gaze changes were edited by Adobe Premiere Pro such that all image content remained the same except for the eye region. Movies consisted of gaze direction changing abruptly once from one frame to another: each dynamic gaze video was composed of 1 frame (40 ms) of the static gaze image at the starting position followed with four repeated frames (160 ms) of the gaze image at the end position. In the left-view faces, gaze could move from the front to the left or in the reverse direction. Similarly, in the right-view faces, gaze could move from the front to the right or in the reverse direction. In frontal faces, gaze could move from the front to either the left or right, or in the corresponding reverse directions. In total there were thus eight combinations of head orientation and gaze shifting patterns for every monkey identity.

We also recorded neural response to static gaze to compare dynamic and static gaze processing in MD. The static gaze stimulus set includes all images that were used to create dynamic gaze shifts. See Yang and Freiwald, 2021 (Yang and Freiwald, 2021) for results of neural tuning to static gaze.

Both dynamic and static gaze stimuli were presented at effective sizes of 16 dva × 12 dva (frontal faces) and 13 dva × 12 dva (left- and right-view faces). Stimuli were shown for 200 ms, followed by 200 ms of grey screen. The midpoint between the two eyes in each gaze was centered on the monitor.

### Data Analysis

#### Neural response

Trials where subject successfully maintained fixation within a time window from 50ms before stimulus onset to 200ms after stimulus offset were included for further analyses. Raw spike trains were first aligned to the stimulus onset of each trial, and then binned using 1 ms bins. Binned spikes trains were averaged across trials to calculate peri-stimulus time histograms (PSTH), which were further smoothed using a Gaussian kernel with standard deviation of 10ms to create the spike density function (SDF).

A cell was classified as responsive to FOB if its SDF to any category of the FOB exceeded 0.8 SD above the baseline for at least 20 consecutive bins (Yang and Freiwald, 2021). The baseline was defined as mean firing rates over a time window from -40 to 10 ms across trials. An FOB response period was then defined for the cell as the combined time periods when the category SDF passed the threshold for all categories the cell was responsive to. If a cell was not FOB-responsive, a default response period from 50 ms to 250 ms was used for FOB. The FOB response period was also used for the dynamic and static gaze experiments, since gaze stimuli were presented in same duration as FOB. For the dynamic faces experiment, a response period lasting from the onset of the FOB response period to 600 ms after was used.

Firing rates were calculated within the response period for each experiment. Normalized neural responses were computed by first subtracting baseline activity and then dividing by the maximum absolute response magnitude.

#### Static face selectivity index

Face selectivity for static images was quantified by a static face selectivity index (SFSI): *SFSI = (R*_*face*_ *- R*_*nonface objects*_*) / (R*_*face*_ *+ R*_*nonface objects*_), in which *R*_*face*_, *R*_*nonface objects*_ denote the mean response (baseline subtracted) to monkey faces and non-face objects respectively. The raw SFSI was adjusted following previous studies (Freiwald and Tsao, 2010): it was set to 1 when *R*_*face*_ > 0 and *R*_*nonface objects*_ < 0, and was set to -1 when *R*_*face*_ < 0 and *R*_*nonface objects*_ > 0. A human static face selectivity index (hSFSI) was calculated using the same formula as the SFSI but with *R*_*face*_ replaced by the response to human faces. Cells with either SFSI or hSFSI larger than 0.33 (i.e. responding at least twice as high to faces than to non-face objects) were classified as static face selective.

#### Dynamic face selectivity index

A dynamic face selectivity index (DFSI) was defined as *DFSI = (R*_*natural*_ *– R*_*object*_*) / (R*_*natural*_ *+ R*_*object*_), where *R*_*natural*_ was the mean response to natural face videos and *R*_*object*_ was the mean response to object videos in the dynamic faces experiment. Both responses were baseline-subtracted, and the same rules for adjusting the SFSI was applied for the DFSI. Cells with DFSI larger than 0.33 were considered dynamic face selective.

#### Face motion selectivity index

A face motion selectivity index (FMSI) was defined as *FMSI = (R*_*natural*_ *- R*_*downsamp5*_*) / (R*_*natural*_ *+ R*_*downsamp5*_), where *R*_*natural*_ was the mean response to the natural face videos and *R*_*downsamp5*_ was the mean response to the 5-frame down-sampled videos, without baseline subtraction.

Wilcoxon rank-sum test was also conducted for each cell to compare its responses to the two conditions. Cells were referred to as face-motion preferred (FM+), when they showed significantly higher responses to the natural face videos (*P* < 0.05, two-tailed), and they were referred to as face-motion suppressed (FM-), when they showed significantly higher responses to the down-sampled videos.

#### Optic flow

Optic flow was estimated for natural face videos using the Farneback method (Farnebäck, 2003) implemented in the Computer Vision Toolbox of MATLAB (Mathworks). Motion direction and magnitude were estimated between frames for each pixel (Fig. 3A).

#### Principal component analysis (PCA) of optic flow and movie-induced neural responses

PCA was performed on the optic flow of the natural face movies and the corresponding population response vectors (Fig. 3B). The horizontal and vertical components of the optic flow were organized into an *N*_*OF*_ *× M*_*OF*_ motion matrix, where *N*_*OF*_, the number of optic flow components, was product of the number of pixels in each frame and the two flow directions (horizontal and vertical); and *M*_*OF*_, the number of video frames with optic flow estimated, was *(N*_*frame*_ *-1) × N*_*video*_. *N*_*frame*_ denoted the number of frames in each video, and *N*_*video*_ was the number of videos. The neural data matrix was created by first horizontally concatenating smoothed PSTHs (0 = 20 ms, time window 0 – 700 ms) to all videos within each FM+ cell and then vertically concatenating data across cells. This led to an *N*_*C*_ *× M*_*C*_ neural matrix, where *N*_*C*_ was the number of cells and *M*_*C*_ was *701 × N*_*video*_. PCA was conducted on the motion and neural matrices separately to find the principle components (PCs). The optic flow and neural data of each movie were then projected onto the PCs individually to create the trajectories in the PC spaces. We employed the first 25 PCs to calculate the Euclidean distance between videos in the space of optic flow or neural responses (Fig. 5C). The first 25 PCs captured 94% of the face optic-flow variance, and 93% of the neural variance. We then performed the same analyses on the optic-flow of object movies and the neural responses to object movies.

PCA was also conducted on the neural responses to gaze stimuli (Figs. 4E and S2). PSTHs of gaze-tuned cells within the time window of 0 – 300 ms were used to build the neural matrix, on which PCA was then performed. Neural trajectories were plotted by projecting neural response to each gaze stimulus onto the first three PCs, which captured 68% of the neural variance.

#### Gaze tuning

Neural tuning to gaze was tested by comparing responses to different gaze conditions within each head orientation by Wilcoxon rank-sum tests (half profile views) or Kruskal-Wallis tests (front views), with the false discovery rate controlled at an alpha level of 0.05.

#### Face tracking and estimation of motion energy

To calculate the motion energy of the head, eyes, and mouth in natural face movies, we first tracked facial features using DeepLabCut (Fig. 5A) (Mathis et al., 2018; Nath et al., 2019). We adopted the primate-face model (Claire Witham, Centre for Macaques, MRC Harwell, UK) provided by DeepLabCut to track the following facial landmarks: “Eyes_MidPoint”, “RightBrow_Top”, “LeftBrow_Top”, “RightEye_Inner”, “LeftEye_Inner”, “RightEye_Outer”, “LeftEye_Outer”, “RightEye_Bottom”, “LeftEye_Bottom”, “OutlineRight_Mouth”, “OutlineLeft_Mouth”, “LowerLip_Centre”, “MidPoint_Nostrils_Mouth”. Regions of interest (ROIs) were then created based on the landmarks tracked on each movie frame. The head ROI was a circle (radius: 2 pixels) centered on the “Eye_MidPoint” landmark. The eye and mouth ROIs were rectangles bounded by the eye and mouth landmarks, respectively (Fig. 5A). We estimated the global motion vector of the head by averaging optic-flow vectors within the head ROI. This global motion vector was then subtracted from the optic-flow vectors within the eye and mouth ROIs to generate head-referenced motion vectors. Motion energy for the head was defined as the magnitude of the global motion vector, while motion energy for the eyes and mouth was defined as the averaged magnitude of the head-referenced motion vectors within the ROIs. Since the optic flow could not be estimated in the first frame of each video, the time series of motion energy started from the second movie frame, with a timestamp at 40 ms. See Movie S3 for the estimated motion energy of all face videos.

#### Modeling the contributions from head, mouth, and eye motion to neural responses: regression analysis of spike trains

Linear regression was conducted to assess the sensitivity of cells to facial motion energy. For each FM+ cell, its spike train was modeled as:

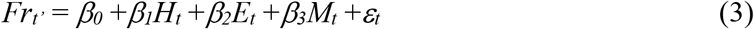

where *t* was the timestamp of each movie frame. *H*_*t*_, *E*_*t*_, and *M*_*t*_ were the motion energy of the head, eyes, and mouth in each frame. To remove multicollinearity among the independent variables, we regressed out *H*_*t*_ from *E*_*t*_ and *M*_*t*_ respectively before adding them into the model (variance inflation factor: 1, 1.06, 1.06 for *H*_*t*_, *E*_*t*_, *M*_*t*_). *Fr*_*t’*_ was the firing rate of the neuron within the time window [*t’, t’*+40], where *t’* = *d* + *t*, with *d* being the delay parameter. Time series of firing rates and motion energy from different videos were concatenated and z-scored before model fitting. To estimate the delay parameter *d*, the spikes trains to the face movies were split into two equal groups of trials. The first group of data was used to optimize the delay parameter *d* by selecting the value leading to a linear model explained the maximum *R*^*2*^ of the neural data. The selected parameter *d* was then adopted in another linear model to fit the second group of data. Neural sensitivity to each motion component was quantified by the standardized *β* coefficients from the model that fit the second group of data.

#### Control analyses for eye motion sensitivity

The first control analysis aimed to confirm that our estimation of the eye motion energy truly reflected local motion of the eyes and was not affected by global head motion. To this end, we created a fake eye ROI by applying the same translation to the original eye ROI across movie frames to move it out of the eye region. This fake eye ROI had the same motion trajectory as the original eye ROI, but did not cover much local face motion within. If our estimation of local motion was accurate, we expected to find local motion energy close to zero within this fake eye ROI. Specifically, the fake eye ROI was created by moving the right eye ROI to the left by 16 pixels (mean distance between the two inner points of the two original eye ROIs). To make sure this fake ROI did not cover any eye region across all frames, its width and length were both shrunk by 50%. Local motion energy within this fake eye ROI was then estimated using the same procedure as that of the true eye ROIs.

The second control analysis was to test if the neural sensitivity to eye motion is specific to a head-referenced frame. We thus created fixed-eye ROIs that did not track the real eyes across movie frames. Two fixed-eye ROIs were created by placing fixed rectangles at the averaged position of the two true eye ROIs respectively. Each fixed-eye ROI has the mean width and length of the corresponding true eye ROI across movie frames. We then estimated the motion energy from the fixed-eye ROI and conducted the regression analysis using the same methods as for the true eye ROIs.

## Acknowledgments

This work was supported by the National Eye Institute of the National Institutes of Health under Award Number (R01 EY021594, W.A.F.), and The New York Stem Cell Foundation. The content is solely the responsibility of the authors and does not necessarily represent the official views of the National Institutes of Health. We thank C. Fisher for help with stimulus preparation; W. Zarco and A. Gonzalez for help with animal training; S. Serene for sharing analysis tools; L. Yin for logistics support; and the veterinary team of the Rockefeller University for the care of the subjects.

## Supplementary Information for

**Fig. S1.**
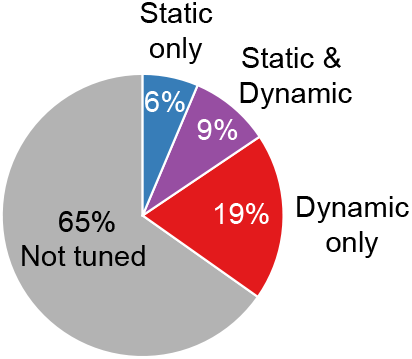
Distribution of face-selective cells tuned to static and dynamic gaze conditions.

**Fig. S2.**
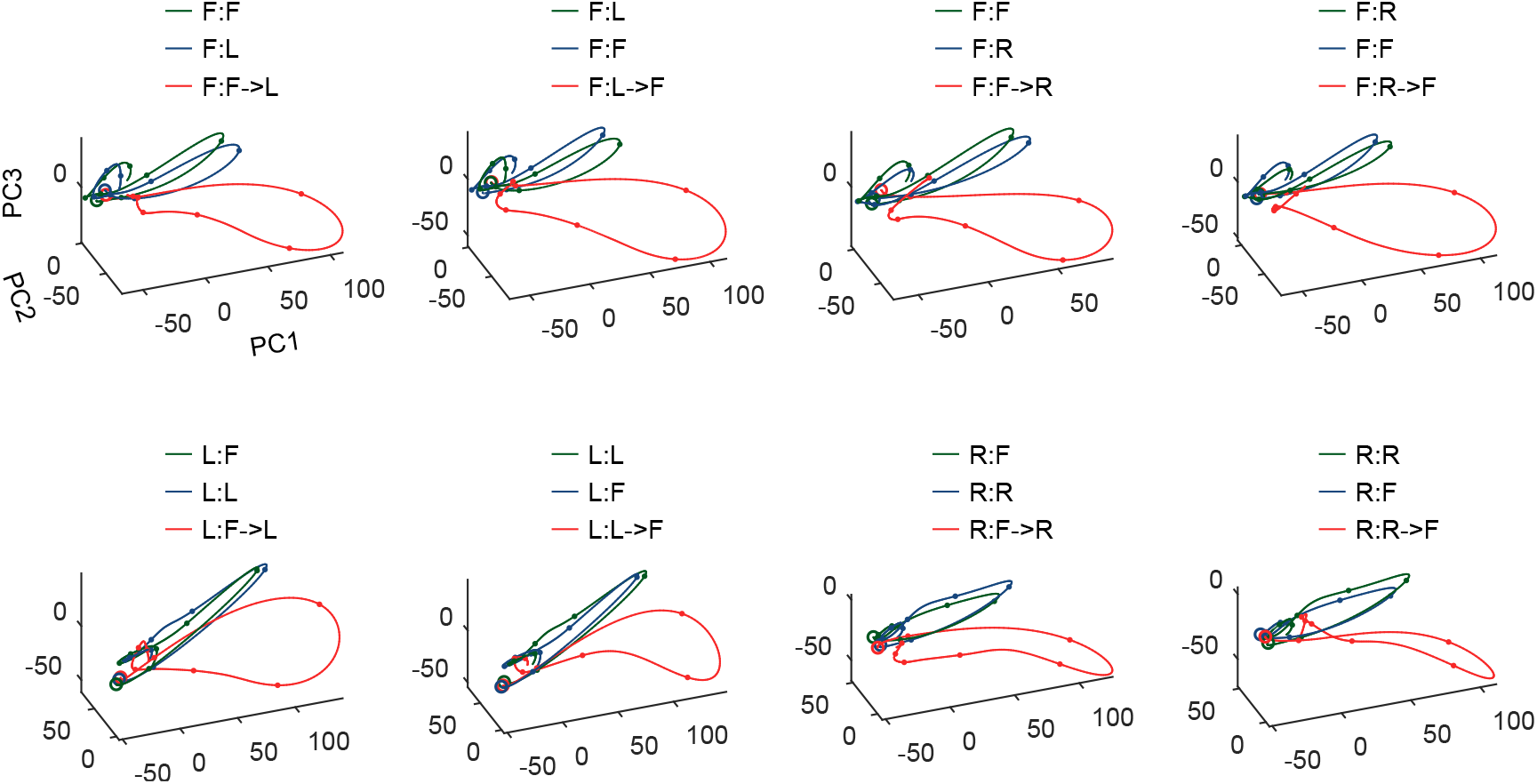
Neural trajectories of gaze-tuned cells in all eight dynamic gaze conditions. For each dynamic gaze condition, trajectories in the two composing static gaze conditions are shown as well. Open circles denote starting states and filled dots indicate 40-ms intervals. The neural trajectories of gaze shifts branched off from those of the static gaze conditions in all eight gaze shifts. Results to one exemplar identity in the stimulus set are shown. Trajectories to other exemplars exhibited similar branching between dynamic and static conditions.

**Fig. S3.**
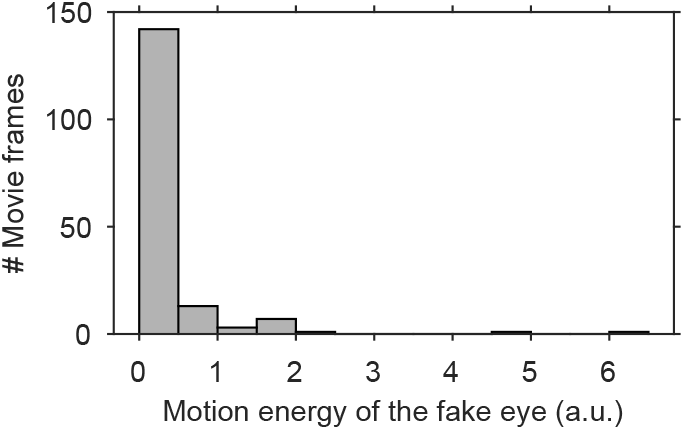
Distribution of motion energy estimated from the fake moving eye across all natural face movie frames.

**Fig. S4.**
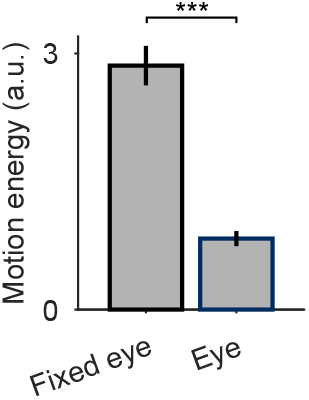
Averaged motion energy of the fixed eyes and the original eyes across natural face movies. Error bars indicate mean ± SEM. \*\*\**P* < 10^−25^ (Wilcoxon signed-rank test).

**Table S1.**
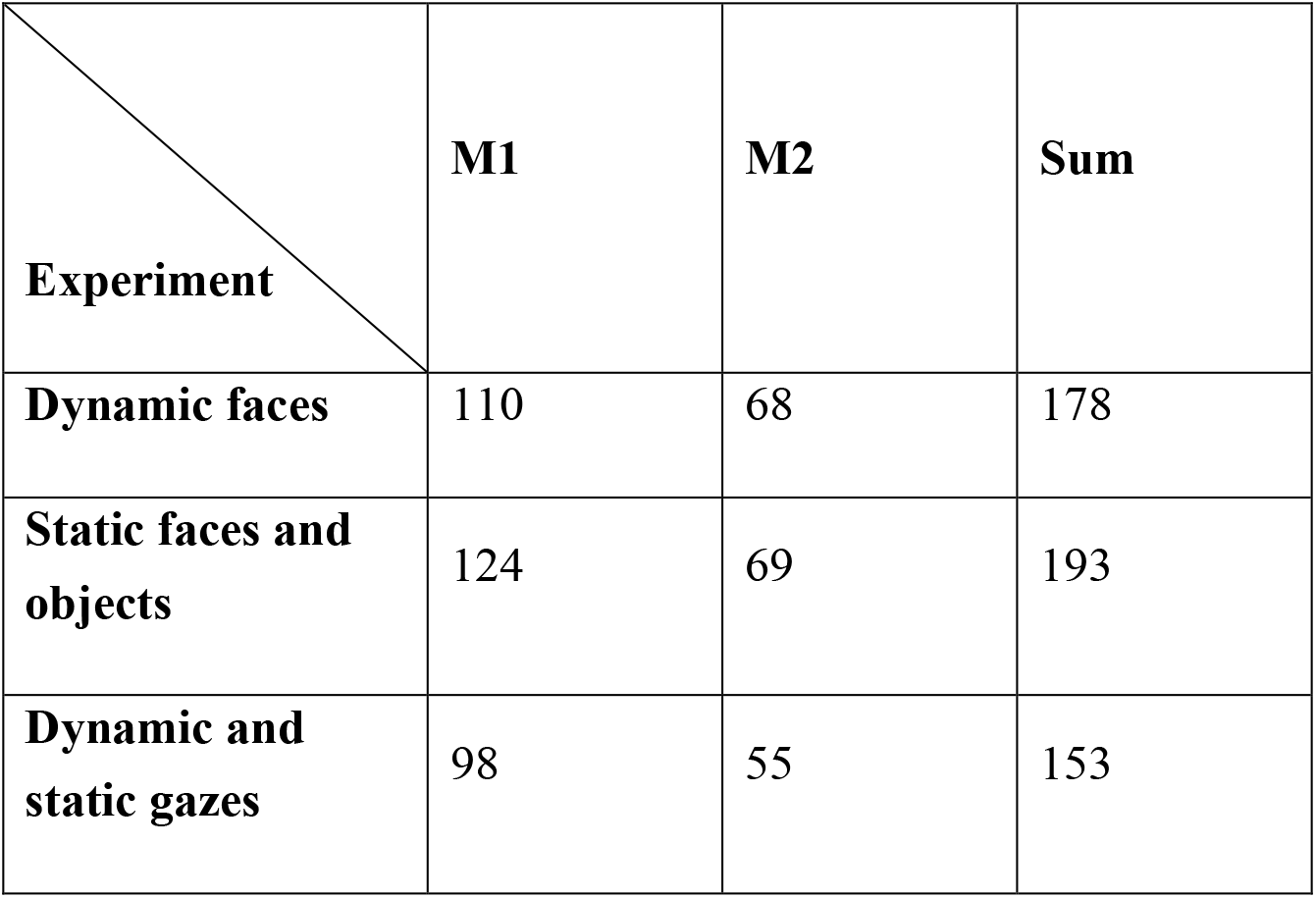
Number of cells recorded in MD from each monkey.

**Movie S1 (separate file)**. Example videos of the eight conditions from the dynamic faces experiment. One exemplar is shown. The whole stimulus set contained 12 exemplars. Texts indicate names of conditions. Frame rate is 25 fps.

**Movie S2 (separate file)**. Example stimuli in the dynamic gaze experiment. One exemplar is shown for demonstration. The whole stimulus set contained three exemplars. Texts indicate labels of conditions. Frame rate is 25 fps.

**Movie S3 (separate file)**. Estimation of facial motion energy. In the video, circles and boxes represent the tracked ROIs for the head, eyes, and mouth. Colors of the ROI borders denote estimated motion energy. No energy was estimated for the first frame of each movie. Results of all 12 natural face movies are shown. Frame rate is 25 fps.

